# A unique serum-free murine cortical astrocyte culture to study endoplasmic reticulum stress in response to amyloid-β

**DOI:** 10.1101/2025.03.10.642375

**Authors:** Diptesh Roy, Sukanya Sarkar, Subhas C. Biswas

## Abstract

Astrocytes are integral to understand Alzheimer’s disease (AD) pathology but the existing serum-supplemented *in vitro* astrocyte culture models are not suitable to study certain stress response mechanisms. Here, we developed a serum-free murine primary cortical astrocyte culture model to study endoplasmic reticulum (ER) stress and inflammation to see the effect of amyloid-beta (Aβ_1-42_). Astrocytes were cultured in a controlled serum-free environment to minimize interference from serum components. Serum-free astrocytes were exposed to oligomeric Aβ to induce ER stress and inflammation. Initially no significant activation of eIF2α, a key marker of ER stress, was observed under serum-free condition but with the removal of N-acetyl cysteine ER stress response was enhanced after 24 hours of Aβ exposure. Subsequently, the inflammatory response, assessed through TNF-α expression, was minimal in the presence of growth factors but became pronounced when these factors were withdrawn. Astrocytic reactivity, assessed by GFAP expression, was observed with prolonged Aβ exposure, indicating a reactive astrogliosis response. Transcript analysis revealed a time-dependent shift in the expression of inflammatory modulators, with early time points showing increased anti-inflammatory markers and late exposure promoting pro-inflammatory responses. These findings highlight the potential of serum-free cultures for studying ER stress and inflammation together in astrocytes and offer insights into the complex role of these cells in AD pathophysiology.

## Introduction

Astrocytes play a vital role in maintaining homeostasis in the central nervous system (CNS) by supporting neurons, regulating blood flow, and managing neurotransmitter levels (1). In Alzheimer’s disease (AD), astrocytes become reactive, promotes neuroinflammation and reduced ability to effectively clear amyloid-beta (Aβ), leading to plaque burden. Their dysfunction further accelerates neurodegeneration by disrupting synaptic function and overall brain health (2-4). The accumulation of misfolded proteins, such as Aβ, induces Endoplasmic reticulum (ER) stress, triggering the unfolded protein response (UPR) in both neurons and glial cells. Prolonged ER stress damages neuronal function by interfering with protein quality control, resulting in synaptic dysfunction and ultimately neuronal death, which speeds up AD progression (5). In astrocytes, ER stress hampers Aβ clearance, may be linked to the neuroinflammatory response worsening AD pathology (6).

Although the blood-brain barrier (BBB) prevents direct access of serum components to brain cells, certain substances like proteins, hormones, and growth factors can still affect the brain. For instance, hormones like insulin and cortisol can enter the brain under physiological conditions, potentially influencing brain activity, mood etc. This is due to the selective permeability of the BBB, which allows specific molecules to cross via active transport or passive diffusion. Beyond those, entry of serum in the brain indicates a possible leakage of BBB under pathological conditions. Since many current *in vitro* astrocyte culture models uses medium supplemented with serum, the main goal of our work is to develop a serum-free astrocyte culture for examining ER stress (7).

Inflammation or changes in the systemic circulation (such as cytokines or other signaling molecules in the blood) can change the permeability of the BBB and cause serum entry, triggering significant effects on the brain cells (8-10). Serum-free media eliminate the variability introduced by serum components (like growth factors, hormones, and proteins), which can otherwise interfere with the correct interpretation of experimental results (11,12).

MD (McCarthy-de Vellis) astrocyte culture containing serum may be useful to address certain biological questions (10) but complicates the interpretation for pathological models including AD. Serum-free primary astrocyte cultures are being developed (11) but the existing protocols are either very expensive (ex. immunopanning) or the addition of several serum-free factors may not suitably allow for the examination of specific biological questions. These limitations highlight the need for a new, effective protocol that allows for a mechanistic analysis of how Aβ accumulation triggers ER stress in cells. Serum-free cultures help better study the direct impact of Aβ on astrocyte function without interference from serum components. In this study, we developed a serum-free murine primary cortical astrocyte culture model to study ER stress and inflammation to see the effect of Aβ.

## Methods

### Materials

Cell culture dishes and flasks were purchased from Nunc (USA). DMEM powder, DMEM (1X), FBS, Trypsin-EDTA, Neurobasal, Sodium pyruvate and L-glutamine were purchased from Gibco (Invitrogen). BSA, Apotransferrin, Putrescine, Progesterone, Selenite, PDL, NAC, HB-EGF, FGF-2 and β-Actin antibody (conjugated with HRP) were the Sigma product. We used eIF2α and Phospho-eIF2α antibody from CST. GFAP antibody were purchased from Novus Biologicals. TNF-α antibody were purchased from Abclonal. The secondary antibody used for immunocytochemistry experiment and western blot were purchased from Invitrogen and CST respectively.

### Primary astrocyte culture

#### A. Isolation of Brain tissue from C57BL/6 mice pups

Before the brain dissection was started, 10 ml of 1X Hank’s Balanced Salt Solution (HBSS) was placed in a 100 mm petri dish, and 10 ml of 0.05% trypsin-BSA solution was prepared. A 0.01% Poly D-Lysine coated T-75 flask was also prepared. Sterile dissection tools, 70% ethanol, ice, and a microscope were arranged for the dissection of mouse pups’ brains. 5-6 mouse pups, born on postnatal day 0 to 1 (P0-P1), were collected. These pups were chosen due to their translucent skin and hairlessness according to Jackson laboratory guide for pup appearance by age. The neck region of the pups was sprayed with 70% ethanol to avoid contamination. The mice pups were decapitated in a single cut using sharp scissors, and the brains were collected into a sterile 10cm dish. The cranium was carefully separated using forceps to expose the brain. Small bent forceps were used to lift the brain and disconnect it from the skull base. The brain was transferred to a sterile 10cm Petri dish containing ice-cold 1X HBSS, ensuring it was fully submerged. The meninges were removed, with careful checks for any residue. Once the meninges were fully removed, the brain was cut into 6–10 pieces using forceps or sharp blades to improve dissociation efficiency. Afterward, the 1X HBSS was aspirated, and 0.05% trypsin-BSA solution was added to the finely chopped cortical tissue, which was then incubated at 37°C for 15 minutes. Following the incubation of the brain pieces, the trypsin-BSA solution was carefully aspirated using a Pasteur pipette. The tissues were transferred to a 15 ml tube with complete astrocyte growth medium (Medium 1), containing DMEM supplemented with 10% fetal bovine serum (FBS) and streptomycin-penicillin (1X).

#### B. Preparation of serum free primary astrocyte culture for treatment

After enzymatic digestion, the tissue was mechanically dissociated into a single-cell suspension. This is typically done by gently triturating or pipetting the tissue to break it into smaller pieces and disperse the cells. The goal is to isolate the astrocytes from the surrounding cells and extracellular matrix while minimizing cell damage. We have to triturate at least 20 times for single cell suspension. Then the cell suspension was poured through a mesh filter (or cell strainer), which allowed smaller individual cells to pass through while capturing larger cell clumps, debris, or tissue fragments. This process created a homogenous single-cell suspension. We have centrifuged the cells in 750 rpm speed for not more than 5mins. Then plated cells into PDL coated T75 flask until 90% confluency in incubator at 37C with 5% CO2. The media is changed every alternate day with medium 1, and when confluency is reached in the mixed cell culture, the flask is gently tapped on both sides several times to remove oligodendrocyte progenitor cells(13). After culturing the cells in Medium 1 for 10 days, they were trypsinized and centrifuged from the T-75 flask. The cells were then plated onto dishes and cover slips with Medium 1 for further use as samples. On the 12^th^ day, two days after subculture, the astrocytes were switched to serum-free Medium 2 (5ng/ml heparin-binding EGF-like growth factor (HBEGF) 50% 1X DMEM (glutamine-less), 50% Neurobasal medium, 1ml L-glutamine (100X), 100μg/ml bovine serum albumin, 100μg/ml apotransferrin, 16 g/ml putrescine dihydrochloride, 60 ng/ml progesterone, 40 ng/ml sodium selenite, 5μg/ml N-acetyl-L-cysteine, 1 mM sodium pyruvate, 100 U/ml penicillin and 100 g/ml streptomycin) (12), deprived of N-acetyl-L-cysteine (NAC) and supplemented with 5 ng/mL of FGF-2 turn it to Medium 3. After maintaining the cells in Medium 3 for 3 days, they were transitioned to Medium 4, a growth factor-deprived medium, on the 15th day. This means that 24 hours prior to treatment, the astrocytes were supplemented with growth factor-deprived media (Medium 4). On the 16th day, the cells were ready for treatment (Fig1).

**Fig. 1.**
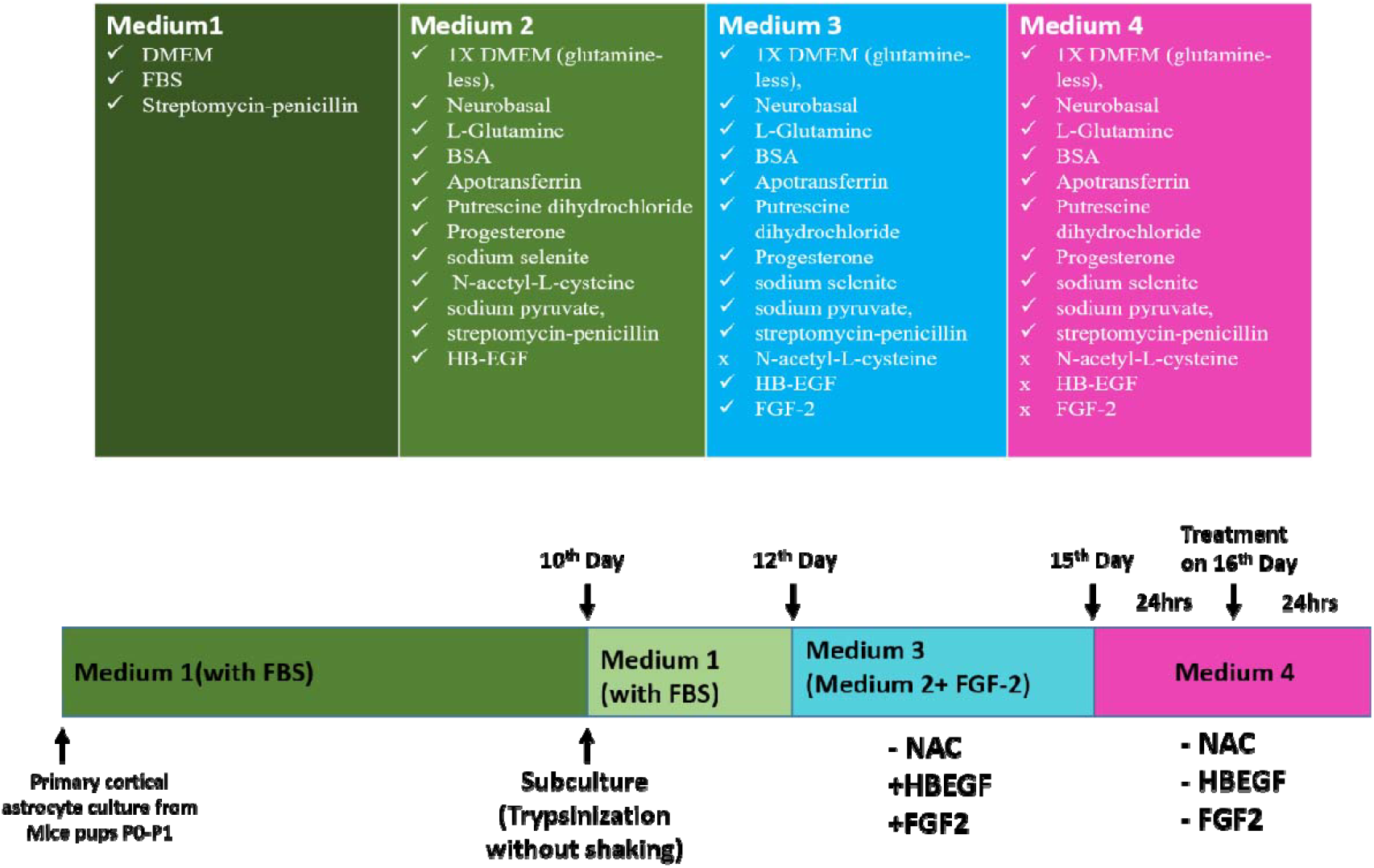
Schematic illustration of serum free culture procedure.

### Oligomeric Aβ preparation

The lyophilized Aβ_1–42_ peptide (American Peptide, Sunnyvale, CA, USA) was first resuspended in 100% 1,1,1,3,3,3 hexafluoro-2-propanol (HFIP) (Sigma-Aldrich) to a concentration of 1 mM and then centrifuged under vacuum conditions in a speed vac (Eppendorf, Hamburg, Germany) until all the HFIP was evaporated. The resulting peptide pellet was dissolved in DMSO (Sigma-Aldrich) at a concentration of 5 mM and subjected to sonication in a 37°C water bath for 10 minutes. The solution was then diluted further with phosphate-buffered saline (PBS: NaCl 137 mM, KCl 2.7 mM, Na2HPO4 10 mM, KH2PO4 2 mM, pH 7.2) and SDS (0.2%) to a final concentration of 400 μM and incubated at 37°C for 6 to 18 hours. Finally, PBS was added to achieve a final concentration of 100 μM, and the solution was incubated for an additional 18 to 24 hours at 37°C. 1.5 μM Aβ_1–42_ was used to treat the mature astrocytes culture at 16^th^ day for different time points.

### Preparation of cell lysate

All treated cells, along with the controls, were washed with PBS and collected by scraping in PBS. The cells were then harvested by centrifugation at 1200 rpm at 4°C for 5 minutes, followed by lysis using lysis buffer (10 mM Tris (pH 4), 150 mM NaCl, 1% Triton X-100, 0.5% NP-40, 1 mM EDTA, 1 mM EGTA, 20 mM NaF, 0.2 mM Orthovanadate, Protease Inhibitors), and incubated on ice for 10 minutes. The cell lysates were then centrifuged at 14,000 rpm at 4°C for 15 minutes. The supernatant was collected, and protein concentration was determined using the Lowry method.

### Western Blotting

Lysates containing equal amounts of protein from each condition were separated using SDS-PAGE. The protein bands were then transferred onto a PVDF membrane. After blocking the membrane with 5% BSA solution for 1.5 hour at room temperature, the membranes were incubated with primary antibodies: anti-mouse β-Actin conjugated with peroxidase (1:10000) for 1 h at room temperature, and anti-rabbit phosphor-eIF2α (1:1000), anti-rabbit total-eIF2α (1:1000), anti-mouse GFAP (1:2500), and anti-rabbit TNF-α (1:1000) antibodies overnight at 4 °C. Following incubation, the membranes were washed three times with TBST [1.5 M NaCl, 1 M Tris (pH 7.5), 0.1% Tween20] and then incubated with HRP-conjugated secondary antibodies for 1–2 h at room temperature, except for the β-Actin blot. Protein bands were detected using the Invitrogen iBright 1500 Imaging System with ECL reagents, following three washes with TBST. Densitometric analysis of the bands was performed using ImageJ software.

### Immunocytochemistry

Both control and treated cells on glass cover slips were fixed with 4% paraformaldehyde in PBS for 10 minutes at room temperature. To permeabilize and block the cells, they were incubated with 3% goat serum and 0.3% Triton X-100 in PBS for 1–2 h. The cells were then incubated overnight at 4 °C with anti-mouse GFAP (1:500). The following day, the cells were washed with PBST (0.3% Triton X-100 in PBS) to remove any excess antibodies. Next, the cells were incubated with species-specific secondary antibodies conjugated to Alexafluor546/488 for 1–2 h at room temperature. Nuclei were stained with Hoechst 33342 at 2 μg/ml in PBS for 30 minutes at room temperature. Images were captured using a LeicaCTR4000 fluorescence microscope with a 40x objective, as indicated in the images. The corrected total cell fluorescence (CTCF) was calculated in ImageJ software, incorporating the integrated density of the staining, the area of the cell, and the background fluorescence from different experimental conditions. CTCF was calculated as: CTCF = Integrated density – (area of selected cell x mean fluorescence of background readings). Data are presented as the mean ± S.E.M. of ten astrocytes from three independent experiments.

Brightfield Microscopic Images were captured without fixation with 20x objective and then adjusted with Photoshop software.

### RNA isolation and Real Time PCR

Total RNA of each sample is isolated from cultured astrocytes by using RNAiso (Takara). From RNA cDNA has been prepared. With cDNA quantitative PCR was performed using TB Green Premix Ex Taq (Tli RNase H Plus) in an Applied Biosystems 7500 Fast Real Time PCR System following the manufacturer’s specifications. Primers used for qPCR; Actin: 5’-GAAATCGTGCGTGACATCAAAG-3’ and 5’-TGTAGTTTCATGGATGCCACAG-3’; Ptx3: 5’-ATGGTTGCTGTGTAGGTGGG-3’ and 5’-GACGACATTTCCCCGGATGT-3’; Sphk1: 5’-GAAACCCCTGTGTAGCCTCC-3’ and 5’-AGCAGGTTCATGGGTGACAG-3’; S100A10: 5’-CAGGTTTGCAGGCGACAAAG-3’ and 5’-CAGCCAGAGGGTCCTTTTGA-3’; C3: 5’-AGCTTCAGGGTCCCAGCTAC-3’ and 5’-GCTGGAATCTTGATGGAGACG-3’; Ggta1: 5’-CCAGTCCCGAGAAGTTCACC-3’ and 5’-CGTTCCTCCAAAAATGGCCG-3’; Iigp1: 5’-GCCACCAATCTTCCTGCTCT-3’ and 5’-CAGCAGCAAATCCTTCCAGC-3’.

### Statistical Analysis

Data are presented as mean ± SEM (standard error of the mean). Statistical significance was determined using one-way ANOVA followed by Tukey’s post hoc analysis. Results were considered statistically significant at p < 0.05.

## Result

### Developing a serum-free media for murine primary cortical astrocyte culture

To develop a serum-free murine primary cortical astrocyte culture for studying ER stress and inflammation in response to Aβ, we employed a serum-free medium that is used earlier. After maintaining the astrocytes in Medium 1 (please see Materials and Methods for details) for 10 days, we observed that the flask was nearly 90% confluent, with astrocytes being the predominant cell type and display a polygonal shape (Fig. 2A). To remove microglia, the cells were subjected to orbital shaking at 100 rpm at 37°C for 8 hours, once pure astrocytes were obtained, the cells were trypsinized for 5 minutes and then centrifuged at 750 rpm for 5 minutes before being subcultured in serum-free Medium 2. After 24 h, the cells began to exhibit signs of deterioration (Fig. 2B), although they still maintained their typical stellate shape. By 72 h, the majority of the astrocytes had died, and only a small number of viable cells remained (Fig. 2C). However, when the cells were replated with Medium 1, following trypsinization and without any shaking, they regained their viability. Microscopic observation over the next two days revealed that, by the 12^th^ day, the flat astrocyte monolayer appeared healthy (Fig. 2D). On the 16th day, after being cultured in Medium 3 for 3 days followed by one day in Medium 4, astrocytes with branched processes emerged (Fig. 2E), mimicking astrocyte’s typical physiological morphology (10).

**Fig. 2:**
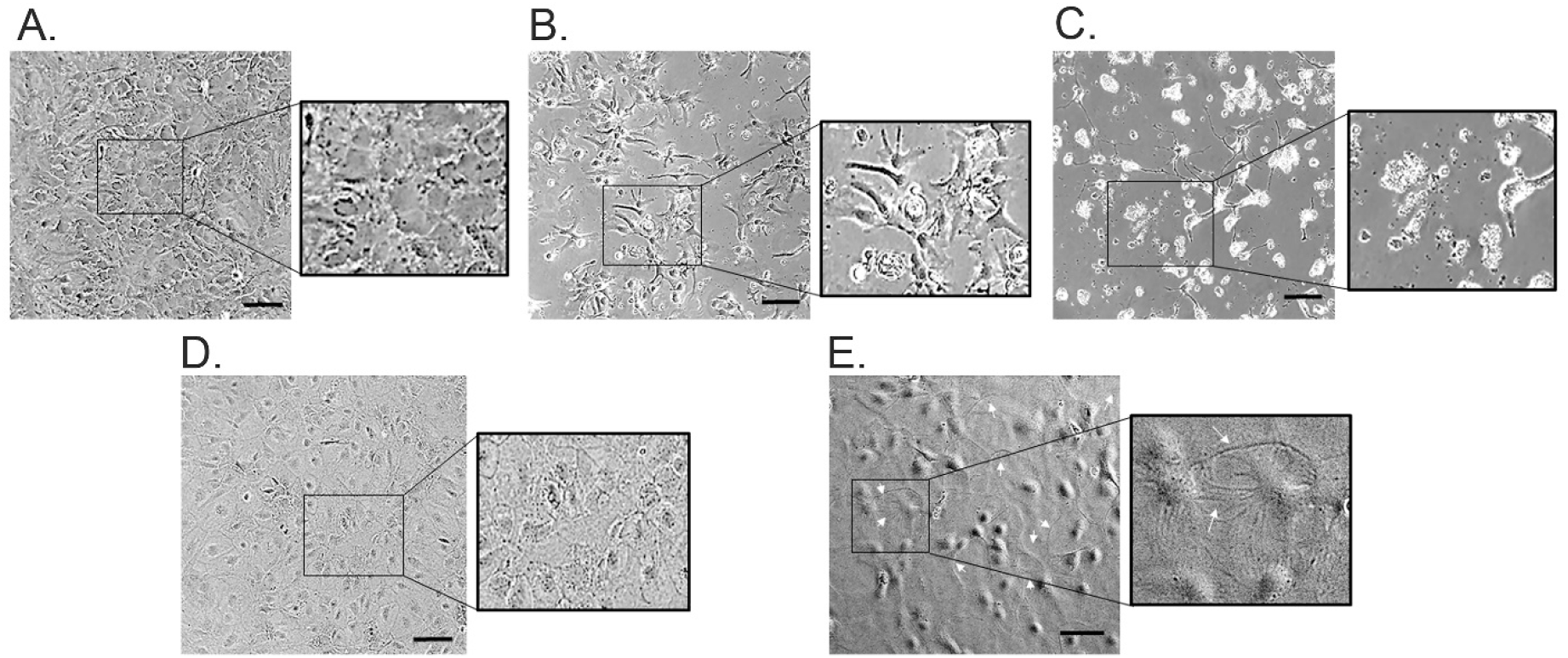
Brightfield microscopic image of astrocyte culture (scale bar 100μ). A. Microscopic image of subcultured astrocytes on 10^th^ day, B. Image of astrocytes after 24h of replate (stellate shape) in Medium 2. C. Image of astrocyte death after 72h of subculture maintained in Medium 2. D. Image of subcultured astrocytes, monolayer of polygonal shaped astrocytes on 12^th^ day in Medium 1, E. Image of astrocytes with branched processes (white arrow) on 16^th^ day.

### Validation of endoplasmic reticulum (ER) stress in astrocytes in response to Aβ

Next, we aimed to assess the impact of Aβ on ER stress response focusing on eIF2α, a key component of the unfolded protein response (UPR) pathway with a particular dosage which is neurotoxic (14). Under conditions of significant ER stress, eIF2α undergoes phosphorylation, which serves as a critical marker of cellular stress (15). After maintaining the astrocytes for 16 days in Medium 2, the cells were treated with 1.5 μM of oligomeric Aβ_1-42_ (henceforth referred as Aβ) at various time points i.e., 0 h, 3 h, 6 h, and 24 h. Western blot analysis was performed to evaluate the phosphorylation of eIF2α. The results showed that irrespective of the duration of treatment (early or late), there was no significant change in the phosphorylation of eIF2α, indicating that Aβ treatment did not induce notable ER stress (Fig. 3 A, B, C) in the astrocytes in Medium 2.

**Fig. 3:**
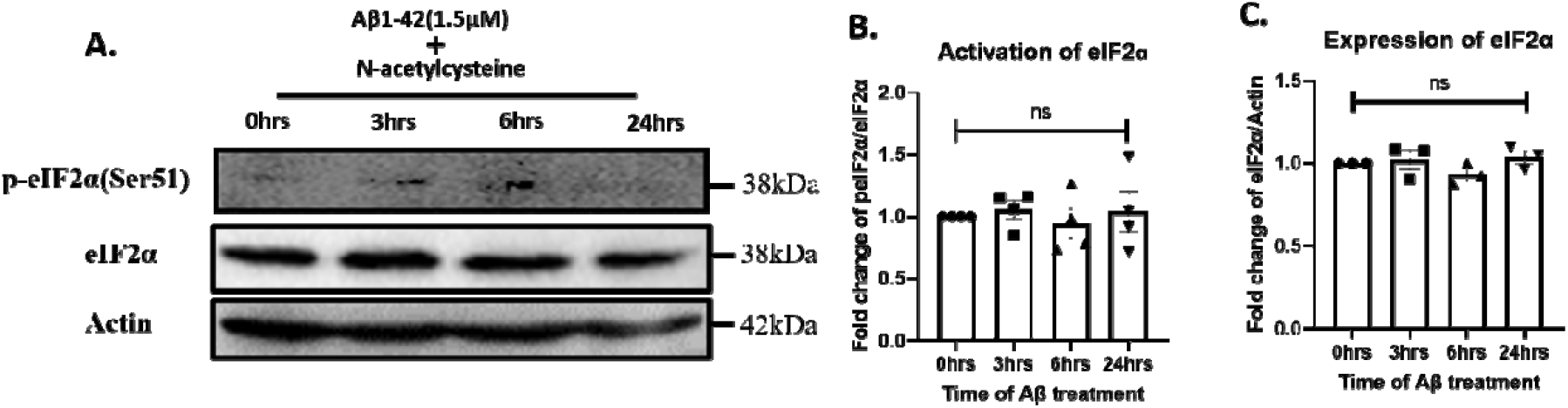
Effect of Aβ on component of UPR. A. Immunoblot of temporal changes in the levels of p-eIF2α, eIF2α & Actin upon Aβ treatment. B, C. Graphical representations of fold change - p-eIF2α/eIF2α (B), eIF2α/Actin (C). Statistical analysis was performed using ANOVA. Data are presented as mean ± SEM (N=3). ns p > 0.05. Actin was used as a loading control.

N-acetyl-L-cysteine (NAC) is a potent antioxidant that gives protection against ER stress (16). Hence, we examined if the removal of NAC from Medium 2 will allow us to study the effect of Aβ on serum-free astrocytes. On the 12th day, the astrocytes were shifted to Medium 3 (without NAC) and as in the previous experiment, Aβ was applied at the same time points (early and late treatment) on day 16. Analysis for the phosphorylation of eIF2α revealed that Aβ induced activation of eIF2α at 24 hours following Aβ treatment, suggesting that ER stress was induced at a later time point compared to the earlier time points. This indicates that prolonged exposure to Aβ (24 h) leads to a more pronounced activation of the UPR pathway when the cells were maintained in Medium 3 (Fig. 4 A, B, C).

**Fig. 4:**
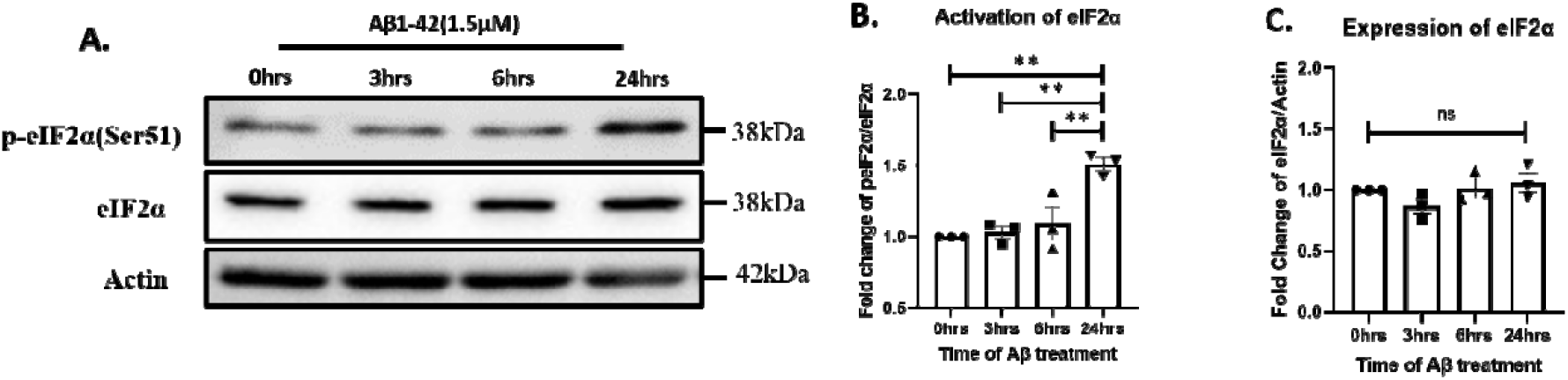
Effect of Aβ on component of UPR when cells are maintained without NAC. A, Immunoblot of temporal changes in expression of p-eIF2α, eIF2α & Actin upon Aβ treatment. B, C, Graphical representations of fold change - p-eIF2α/eIF2α (B), eIF2α/Actin (C). Statistical analysis was performed using ANOVA. Data are presented as mean ± SEM (N=3). ns p > 0.05, **p < 0.01. Actin was used as a loading control.

We next examined if Medium 3 will allow us to explore the effect of Aβ on inflammation and ER stress together in this astrocyte model since inflammation and ER stress are closely related in the context of AD. We measured the protein expression of TNF-α, a key inflammatory cytokine. However, there were no significant change in TNF-α expression at any of the time points (0 h, 3 h, 6 h, and 24 h) after Aβ treatment to primary astrocytes on 16^th^ day in Medium 3 (Fig.5 A, B).

**Fig. 5:**
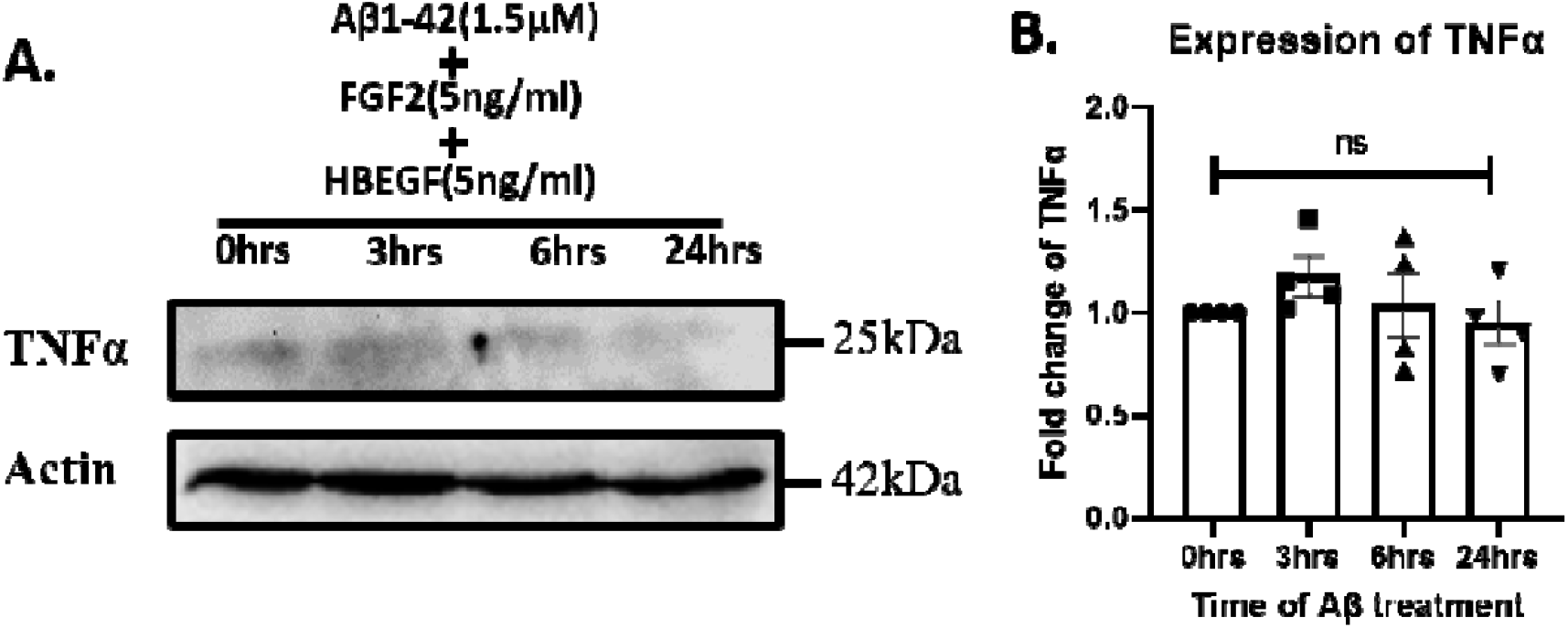
Effect of Aβ on TNF-α. A. Immunoblot of temporal changes in expression of TNF-α & Actin upon Aβ treatment. B. Graphical representation of fold change TNF-α/Actin. Statistical analysis was performed using ANOVA. Data are presented as mean ± SEM (N=3). ns p > 0.05. Actin was used as a loading control.

Presence of specific growth factors can sometimes mask the effect of Aβ on astrocytes (17). Hence, on the 16th day, we exposed the cells to Aβ at four different time points (0, 3, 6, and 24 h) to evaluate inflammation in astrocytes in the absence of growth factors FGF-2 & HBEGF (Medium 4). We found that the expression of TNF-α was significantly elevated following 24 h of Aβ treatment (Fig. 6 A, B).

**Fig. 6:**
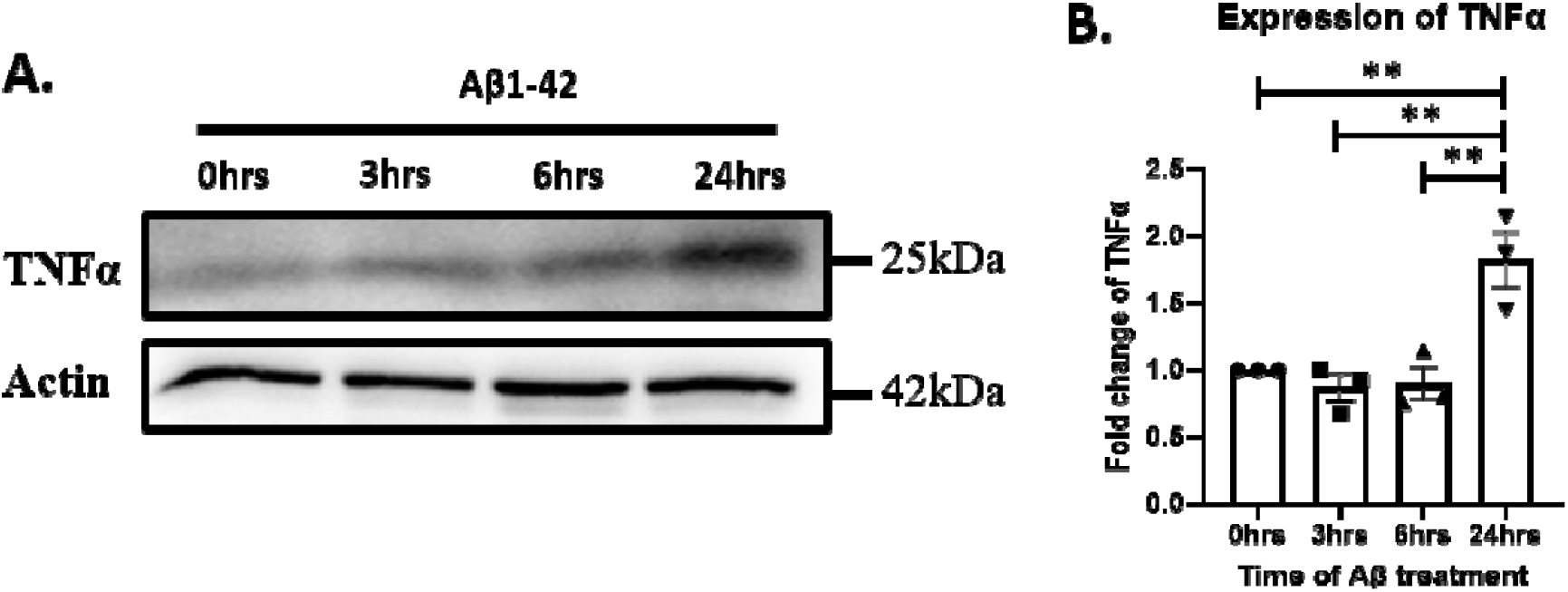
Effect of Aβ on TNF-α when FGF-2 & HBEGF has been withdrawn from medium. A. Immunoblot of temporal changes in expression of TNF-α & Actin upon Aβ treatment. B. Graphical representation of fold change TNF-α/Actin. Statistical analysis was performed using ANOVA. Data are presented as mean ± SEM (N=3). ns p > 0.05, **p < 0.01. Actin was used as a loading control.

### Astrogliosis in response to Aβ treatment

Further, we examined how Medium 4 changes the reactivity of astrocytes in the presence of Aβ at the same time points as earlier to evaluate the morphological shift and changes in expression of GFAP, an astrocyte marker. Using western blot analysis, we found that GFAP was increased significantly after 24 h exposure of Aβ compared to the earlier time points (Fig. 7 A, B). Immunofluorescence analysis of GFAP expression revealed that upon 24 h of Aβ exposure the CTCF (corrected total cell fluorescence) of GFAP was increased in astrocytes in Medium 4 (Fig. 7 C, D) versus the earlier time points. Moreover, cortical astrocytes were transformed into more fibrous shape after 24 h treatment assessed by individual cell perimeter which was significantly increased compared to the earlier time points (Fig. 7E).

**Fig. 7:**
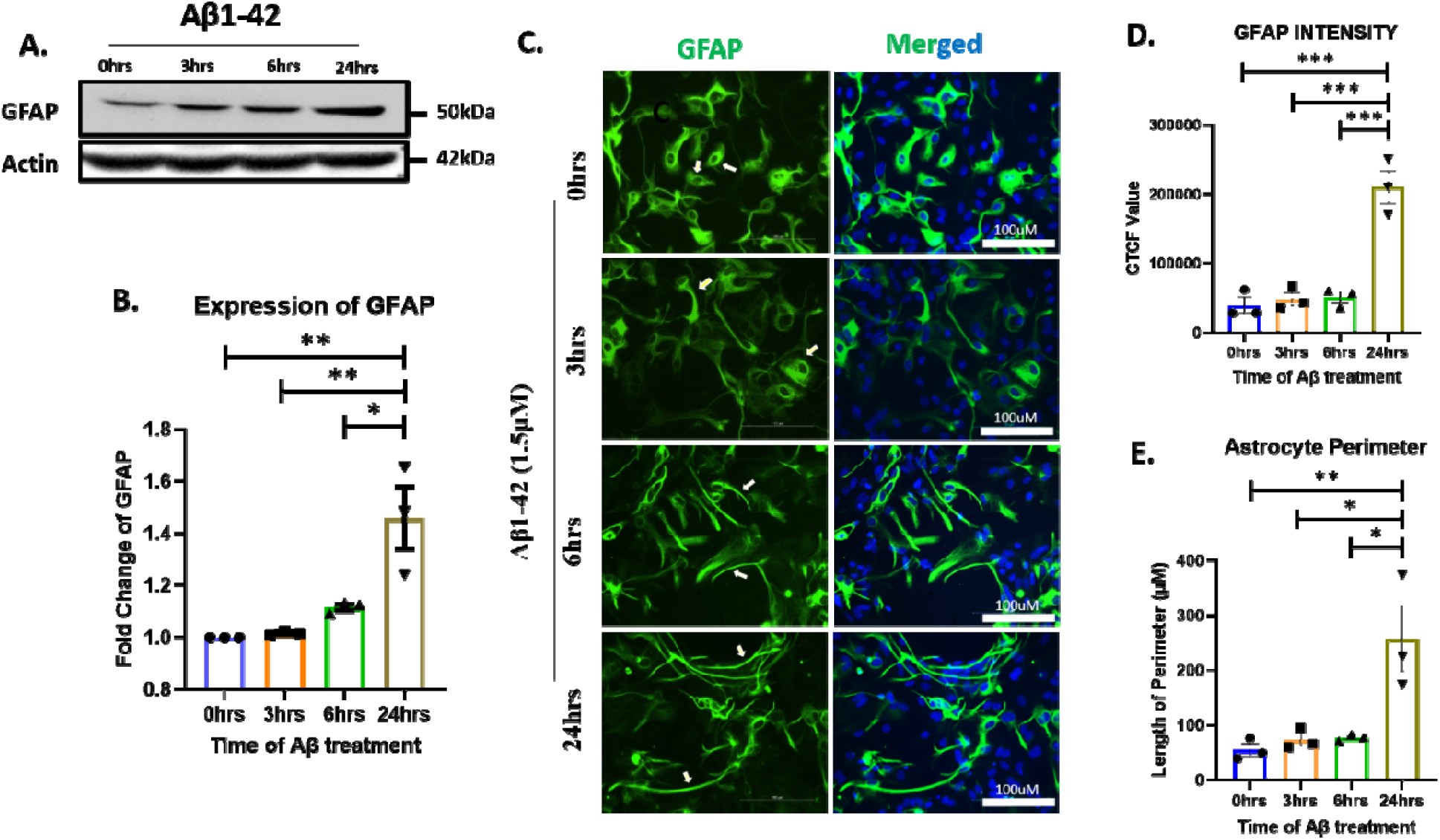
Astrocyte reactivity upon Aβ treatment. A. Immunoblot of GFAP & Actin of primary astrocyte treated with oligomeric Aβ, B. Graphical analysis of fold change of GFAP/Actin, C. Images of primary astrocyte treated with oligomeric Aβ. Immunocytochemistry was done using anti-GFAP (green) antibody & DAPI (blue) for nucleus, D. Graphical representation of corrected total cell fluorescence of GFAP and E. length of astrocytes perimeter treated with Aβ. Statistical analysis was performed using ANOVA. Data are presented as mean ± SEM (N=3). *p < 0.05, **p < 0.01, ***p < 0.001. Actin was used as a loading control.

### Characterization of astrocyte-specific inflammatory molecules upon Aβ treatment

Finally, we characterized these Medium 4 astrocyte-rich culture for astrocyte-specific inflammatory molecules. Accordingly, the Aβ treated samples (at different time points - 0, 3, 6, and 24 h) were collected to examine the gene expression levels of several inflammatory molecules (9) and found that the transcript levels of anti-inflammatory molecules like Ptx3, S100a10 and Sphk1 (18) were increased (Fig.8 D, E, F) at early time points whereas the pro-inflammatory molecules like C3, Iigp1, Ggta1 (19) were increased at 24 h of Aβ exposure (Fig. 8 A. B, C).

**Fig. 8:**
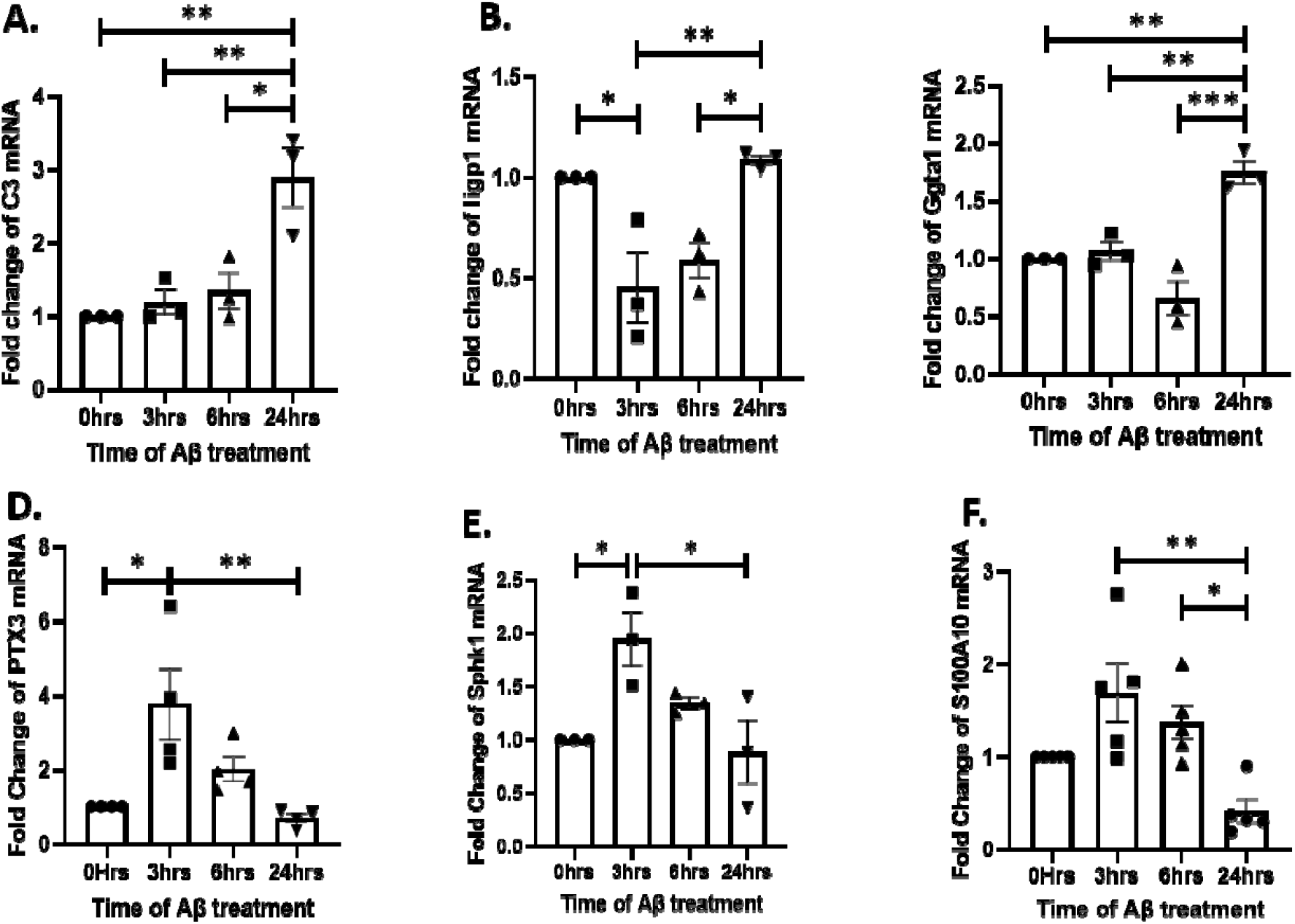
Characteristics of inflammatory molecules in serum-free astrocytes upon Aβ treatment by RT-PCR. Graphical representation of fold change of A. C3, B. Iigp1, C. Ggta1, D. PTX3, E. Sphk1, F. S100A10. Statistical analysis was performed using ANOVA. Data are presented as mean ± SEM (N=3). *p < 0.05, **p < 0.01, ***p < 0.001. Actin was used as a loading control.

## Discussion

ER stress and inflammatory pathways are implicated in AD pathogenesis (15,20) but how subtle ER stress leads to inflammatory responses remains elusive. Alongside neurons, studying ER stress and inflammatory responses in astrocytes are critical for integrative mechanistic understanding of these pathways in AD. Although there are well-established methods for culturing neurons to study ER stress responses *in vitro*, current literature lacks in cell culture protocols dedicated for studying ER stress responses in astrocytes. We aimed to study ER stress in primary cortical astrocytes in response to Aβ. After observing the lack of response to Aβ treatment in serum-free primary cultures of astrocytes using a protocol reported earlier (10), we identified a compound, NAC, which appears to have an inhibitory effect on ER stress (16). NAC is a potent antioxidant that has been shown to have protective effects against various forms of cellular stress, including ER stress (21). NAC functions primarily by replenishing intracellular levels of glutathione, a key antioxidant in the cell. By enhancing glutathione levels, NAC reduces oxidative stress, which is a significant contributor to ER stress. In the context of ER stress, NAC has been reported to alleviate cellular damage caused by the accumulation of misfolded proteins in the ER lumen, a key feature of the unfolded protein response (UPR). It can mitigate oxidative stress, promote protein folding, and restore normal cellular functions, thus potentially protecting cells from the harmful effects of prolonged ER stress. We also found that growth factors like FGF-2 and HBEGF play a protective role against pro-inflammatory cytokines (22,23). To explore this further, we excluded these components from the modified medium (which did not contain NAC) and on the 15^th^ day (24 h prior to Aβ treatment) the cells were cultured in modified medium without FGF-2 and HBEGF (referred to as Medium 4). FGF-2 has been shown to regulate inflammatory responses by reducing the production of pro-inflammatory cytokines (24), while HBEGF helps to reduce the secretion of these mediators. Furthermore, HBEGF can act as a protective factor under inflammatory conditions by promoting cell survival and preventing the overall inflammatory response. For the experiment, the cells were maintained in Medium 4 on the 16th and 17th days, so that the experiment was conducted without the inclusion of growth factors to eliminate their potential inhibitory effects on inflammation ensuring that even subtle effects of Aβ exposure could be quantified, which further underscores the impact of these growth factors in modulating the inflammatory response in astrocytes (24,25). The use of the specific medium for culturing astrocytes in this study is critical for maintaining cell viability, promoting growth, and preserving the appropriate cellular responses such as ER stress. In summary, the chosen medium provides the necessary conditions to sustain astrocytes in an optimal state for both maintenance and experimental treatment, removing confounding factors in interpreting results in studies of ER stress, inflammation and Aβ effects.

The results of astrogliosis study have important implications for understanding the role of astrocytes in the context of AD and Aβ toxicity. The significant increase in GFAP expression suggests that astrocytes undergo a reactive response to Aβ exposure. This is consistent with the concept of reactive astrogliosis, where astrocytes become reactive and upregulate GFAP expressions in response to neuroinflammatory stimuli (26-28). The evaluation of characteristics of astrocytes by transcripts profile indicate a time-dependent shift in the expression of inflammatory modulators in astrocytes exposed to Aβ. Early time points (0, 3, 6 h) showed increased transcript levels of anti-inflammatory modulators such as Ptx3, S100a10, and Sphk1, suggesting an initial protective or regulatory response. However, after 24 h of Aβ exposure, pro-inflammatory modulators like C3, Iigp1, and Ggta1 were significantly upregulated, indicating a transition toward a more inflammatory state in astrocytes (9). These findings have important implications for understanding the dynamic role of astrocytes in AD. The early anti-inflammatory response could be an attempt by astrocytes to counteract Aβ-induced damage, but the prolonged exposure appears to push astrocytes toward a pro-inflammatory phenotype, which may contribute to neurodegeneration (18,29,30).

## Conclusion

In conclusion, the serum-free culture conditions established in this study provide a controlled and reproducible model for studying astrocyte responses to Aβ. The results demonstrate that astrocytes exhibit a time-dependent shift from anti-inflammatory to pro-inflammatory responses, highlighting their complex role in neuroinflammation and neurodegeneration. Hence, this unique culture medium, which supports astrocyte health and function, is crucial for investigating ER stress and inflammation pathways together in AD.

## Abbreviations

Aβ: amyloid-beta
AD: Alzheimer’s Disease
ER: Endoplasmic reticulum
eIF2α: Eukaryotic Initiation Factor 2α
TNF-α: Tumor Nectrosis Factor α
GFAP: Glial fibrillary acidic protein
NAC: N-acetyl-L-cysteine
FGF-2: Fibroblast Growth Factor-2
HBEGF: Heparin Binding EGF-like growth factor
BBB: Blood-Brain Barrier

## Acknowledgement

We would like to thank Mr. Sounak Bhattacharya from the Central Instrument Facility (CIF), CSIR-IICB, Kolkata for his contributions to carry out Leica Sp8 STED confocal microscopy image acquisition. We thank Dr. Priyankar Sanphui for valuable discussions. We thank Dr. Debabrata Biswas for providing psPAX2 and pMD2 plasmids for lentivirus preparation.

## Author contributions

D.R. and S.C.B conceived and designed the study. S.C.B. supervised the study, acquired funding for the experiments and provided all the necessary resources for carrying out the work. D.R. did all the biochemical and immunofluorescence studies. D.R and S.S. designed and carried out the RT-PCR studies. D.R. analysed all the data and wrote the original draft of the manuscript. S.S. and S.C.B. reviewed and edited the manuscript and contributed to writing of the paper.

## Funding

This work was supported partly by CSIR-Indian Institute of Chemical Biology, Govt. of India; Grant number: OLP-115.

## Competing interest

No potential conflict of interest was reported by the author (s).

## Ethical approval

All animal studies were carried out in accordance with the guidelines formulated by the Committee for Control and Supervision of Experiments on Animals (Ministry of Fisheries, Animal Husbandry and Dairying, Department of Animal Husbandry and Dairying, Govt. of India) with approval from the Institutional Animal Ethics Committee (IAEC). We have consulted the ARRIVE guidelines for the relevant aspects of animal studies.

## Notes

### Competing Interest Statement

The authors have declared no competing interest.

